# Differences in single-motor and multi-motor motility properties across the kinesin-6 family

**DOI:** 10.1101/2022.07.13.499883

**Authors:** Andrew Poulos, Breane G. Budaitis, Kristen J. Verhey

## Abstract

Kinesin motor proteins are responsible for orchestrating a variety of microtubule-based processes including intracellular transport, cell division, cytoskeletal organization, and cilium function. During cell division, members of the kinesin-6 family play critical roles in anaphase and cytokinesis, however little is known about their motility properties. We find that truncated versions of MKLP1 (*Hs*KIF23), MKLP2 (*Hs*KIF20A), and *Hs*KIF20B predominately display non-processive behavior as single molecules although slow, processive motility was occasionally observed, most prominently for MKLP2. Despite their non-processive nature, all kinesin-6 proteins were able to work in teams to drive microtubule gliding. MKLP1 and KIF20B were also able to work in teams to drive robust transport of both peroxisomes, a low-load cargo, and Golgi, a high-load cargo, in cells. In contrast, MKLP2 showed minimal transport of peroxisomes and was unable to drive Golgi dispersion. These results indicate that while all three mammalian kinesin-6 motor proteins are generally non-processive as single motors, they differ in their ability to work in teams and generate forces needed to drive cargo transport in cells.

## Introduction

Kinesins are a superfamily of proteins responsible for orchestrating fundamental microtubule-based processes including cell division, intracellular trafficking, cytoskeletal organization, and cilium function [1-4]. Kinesin proteins are defined by a highly-conserved kinesin motor domain that contains signature sequences for nucleotide and microtubule binding. Sequence differences within this motor domain result in motor-specific motility properties that are tuned to the functional output of that motor in cells. The kinesin-6 family is best known for its roles in mitosis and cytokinesis in animal cells [5-8] although these proteins also play important contributions in interphase of cycling cells as well as in post-mitotic cells [9, 10]. The kinesin-6 family consists of three subfamilies, two of which are conserved across eukaryotes, mitotic kinesin-like protein 1 (MKLP1: *Hs*KIF23, *Dm*Pavarotti, *Ce*ZEN-4) and mitotic kinesin-like protein 2 (MKLP2: *Hs*KIF20A/RAB6KIFL, *Dm*Subito), whereas the third subfamily is vertebrate-specific, KIF20B [also known as mitotic phosphoprotein 1 (MPP1)] (Figure 1A).

**Figure 1.**
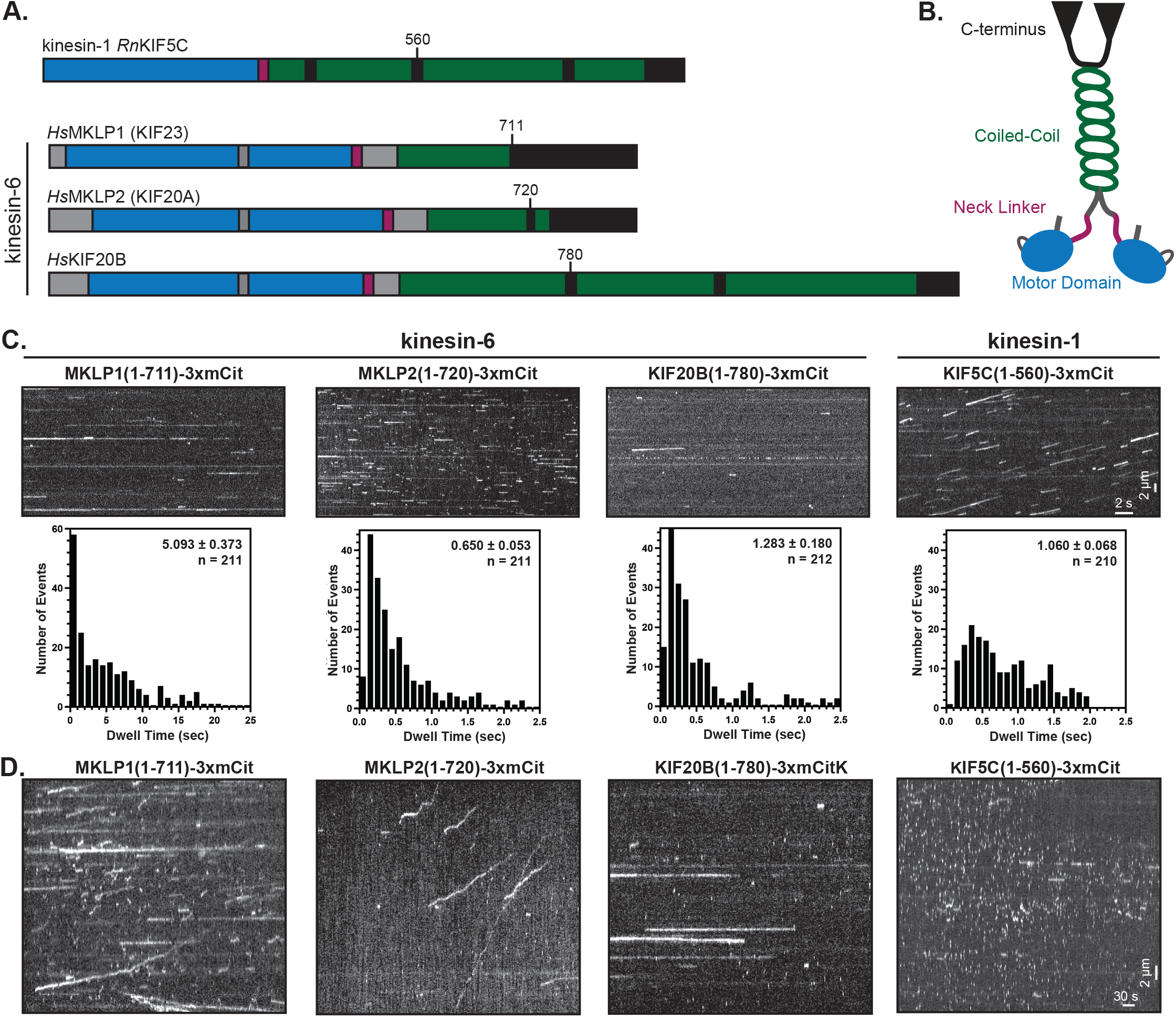
Single-molecule motility properties of kinesin-6 motors. (A) Domain organization and (B) schematic of kinesin-6 motor proteins. Blue, motor domain; green, coiled-coil; purple, neck linker; gray, insertions in kinesin-6 relative to kinesin-1. Numbers and black lines indicate the positions of protein truncations. (C) Motility properties of the indicated kinesin-6 proteins and the kinesin-1 control using a fast acquisition rate (1 frame/50 ms). Representative kymographs are shown with time on the x-axis (scale bar, 2s) and distance on y-axis (scale bar, 2 μm). Each motor’s dwell time on the microtubule was calculated from kymographs and plotted as a histogram for the population. Insets display the mean ± SEM and number of events across three independent experiments. (D) Motility properties of indicated kinesin-6 proteins and the kinesin-1 control using a slow acquisition rate (1 frame/2 s). Representative kymographs are shown with time on the x-axis (scale bar, 30s) and distance on the y-axis (scale bar, 2 μm).

During anaphase, the central region of the mitotic spindle is remodeled to form a narrow region of antiparallel, overlapping microtubules called the central spindle or midzone. Formation of the central spindle requires the function of MKLP1 and MKLP2. MKLP1 is a component of the centralspindlin complex [11-13], which accumulates at the central spindle and promotes anti-parallel microtubule bundling during central spindle assembly [12, 14-19]. MKLP2 also accumulates at the central spindle and is responsible for relocalization of the Chromosome Passenger Complex (CPC) components, including Aurora B kinase, from the centromeres to the central spindle [20-29]. During telophase, the central spindle is further remodeled to form the midbody which connects the two daughter cells into G1. During cytokinesis, MKLP1 and MKLP2 also play critical roles in the assembly and constriction of the contractile ring, an assembly of unbranched actin filaments and myosin-II, by directing the formation of a zone of active RhoA at the cortex of the cell equator [26, 30-34]. Inhibition of centralspindlin activity or MKLP2-driven delivery of the CPC leads to a reduction in constriction rate and increase in cytokinesis failure [11, 12, 17, 24, 27, 35-48]. Much less is known about the cellular functions of KIF20B which regulates midbody maturation and is necessary for the completion of cytokinesis [49-51].

How sequence divergence of the kinesin motor domain gives rise to the evolution of distinct motility properties is critical to understanding the functional roles played by specific kinesins in cells. Many kinesin proteins utilize stepwise processivity as their mode of motility and thereby drive microtubule-based transport of cargo. However, the motility properties of members of the kinesin-6 subfamilies are largely unknown. The kinsesin-6 family is defined by three features of the motor domain: an N-terminal extension, an insertion in loop-6 on the surface of the motor domain, and an extended region between the neck linker and the first coiled coil predicted to drive homodimerization [12, 51-54] (Figure S1C). The neck linker connects the catalytic motor domain to the coiled-coil domain and is important in force production and mechanochemical coordination of the two motor domains in a dimeric kinesin protein [55, 56]. The extended neck linker of kinesin-6 proteins suggests that kinesin-6 proteins may not perform classical stepwise movement [12, 18, 27, 53, 57].

The motility properties of individual kinesin proteins are typically examined in single-molecule motility assays where processive motors show unidirectional movement over time and non-processive motors show either diffusive or static binding along the microtubule lattice. However, an inability to undergo processive motility does not preclude a kinesin from driving microtubule-based transport events as non-processive kinesin proteins can drive motility [58-61]. Single-molecule motility behavior of mammalian MKLP1 has not been evaluated, but this motor can drive the movement of axonemes (microtubule bundles) in a gliding assay [14]. More progress has been made with the *C. elegans* homolog, *Ce*ZEN-4, which undergoes diffusive movement along the microtubule lattice as single motors, can undergo processive motility as clusters of motors, and can drive motility in microtubule gliding assays [18, 62]. *Ce*ZEN-4 and *Dm*Pav have also been observed to bundle microtubules in *in vitro* assays [12, 42, 62-64] and have thus been suggested to function in the bundling of antiparallel microtubules at the central spindle.

For MKLP2, recent research showed that the full-length protein undergoes processive motility as single molecules and can transport purified CPC complexes along microtubules [29], suggesting that MKLP2 may be capable of a classic kinesin transport function. However, when imaging MKLP2 and CPC components in dividing cells, the majority of MKLP2-CPC particles displayed diffusive behavior and only a small number of events showed behavior consistent with directed movement [27, 29]. For *Hs*KIF20B, its single-molecule motility properties have not been described but it is capable of multi-motor transport in a microtubule gliding assay [49]. Finally, the sole member of the kinesin-6 family in fission yeast, *Sp*Klp9, forms homotetramers that display slow plus end-directed motility both as single molecules and in microtubule-gliding assays [65].

We utilized a variety of assays to characterize the single-molecule and multi-motor motility properties of kinesin-6 motors MKLP1, MKLP2, and KIF20B to understand how their motility properties relate to their functions in cells. We find that generally, kinesin-6 motors are not capable of directed motion as single motors but can work in teams to drive transport in gliding assays and cargo-driven transport in cells. Our results provide a basis for understanding kinesin-6 motility properties and can provide insight into how mutations in kinesin-6 motors can lead to disruption of neural development or cancer [66-68].

## Results

### Kinesin-6 proteins are non-processive as single motors

We first tested whether kinesin-6 motors could undergo processive motility as individual motors using a standard single-molecule motility assay. Because many kinesin proteins utilize their C-terminal tail domains for autoinhibiton of the N-terminal motor domain [64, 69, 70] and/or as an auxiliary microtubule-binding domain [71-76], we generated truncated versions of each kinesin-6 protein that contain the kinesin motor domain and a portion of the predicted coiled-coil segment for dimerization (Figure S1A). For MKLP1 (*Hs*KIF23), we compared the truncated MKLP1(1-711) protein to the full-length MKLP1(1-856). For MKLP2 (*Hs*KIF23), we tested two truncated versions, MKLP2(1-720) and MLKP2(1-770) and for KIF20B, we also tested two truncated versions, KIF20B(1-780) and KIF20B(1-982). All motors were tagged at their C-terminus with three tandem mCitrine (3xmCit) fluorescent proteins for fluorescence imaging. In preliminary experiments, the shorter and longer versions of each protein behaved similar to each other (Figures 1 and S2), therefore only the results from the shorter truncations will be described in detail.

We used single-molecule imaging to examine the ability of the truncated motors to undergo processive motility along taxol-stabilized microtubules. The well-characterized kinesin-1 protein KIF5C(1-560)-3xmCit [77] was used as control (Figure 1A). Under standard imaging conditions (1 frame/50 ms), all kinesin-6 motor proteins were observed to transiently bind to and release from microtubules without undergoing directional motility with average dwell times of 5.093 ± 0.373 sec for MKLP1(1-711), 0.650 ± 0.053 sec for MKLP2(1-720), and 1.283 ± 0.180 sec for KIF20B(1-780) (Figure 1C). All motors displayed similar behaviors when tagged with Halo and Flag tags (Figure S3), suggesting that the 3xmCit tag did not cause a detrimental effect on motility.

It is possible that kinesin-6 motors undergo slow processive motility that can be difficult to observe using standard imaging conditions. To test this, we repeated the motility assays at a slower imaging rate (1 frame/2 s). Most molecules again underwent transient binding and release from the microtubules, however, some processive, unidirectional events were observed for each kinesin-6 motor (Figure 1D). Processive motility events were most frequently observed for MKLP2(1-720)-3xmCit [23% of events (n = 200 total)] whereas fewer processive events were observed for MKLP1(1-711)-3xmCit [14% of events (n = 150 total)] and KIF20B(1-780)-3xmCit [1.3% of events (n = 150 total)]. When moving processively, all kinesin-6 motors showed inconsistent walking behavior, with frequent pausing and varying velocities. Taken together, the single-molecule assays show that kinesin-6 motors are largely non-processive as individual motors and may need to work in teams to drive transport.

### Kinesin-6 proteins can work in teams to drive microtubule gliding

We used a microtubule gliding assay to test whether kinesin-6 motors can work in teams to drive motility. To do this, kinesin-6 motor proteins were biotinylated by fusion of an AviTag to their C-termini and co-expression with the enzyme BirA. The biotinylated motors were statically attached to a neutravidin-coated coverslip and then taxol-stabilized microtubules were introduced into the chamber (Figure 2A). Interestingly, all three kinesin-6 motors were able to work in teams to glide microtubules (Figure 2B). In each case, the speed of microtubule gliding was slow, similar to the speeds observed in single-molecule motility assays (Figure 1), with average velocities of 43.9 nm/s for MKLP1, 60.6 nm/s for MKLP2, and 61.7 nm/s for KIF20B. These results indicate that although the kinesin-6 motor proteins are largely non-processive as single molecules, they can work in teams to drive slow microtubule gliding.

**Figure 2.**
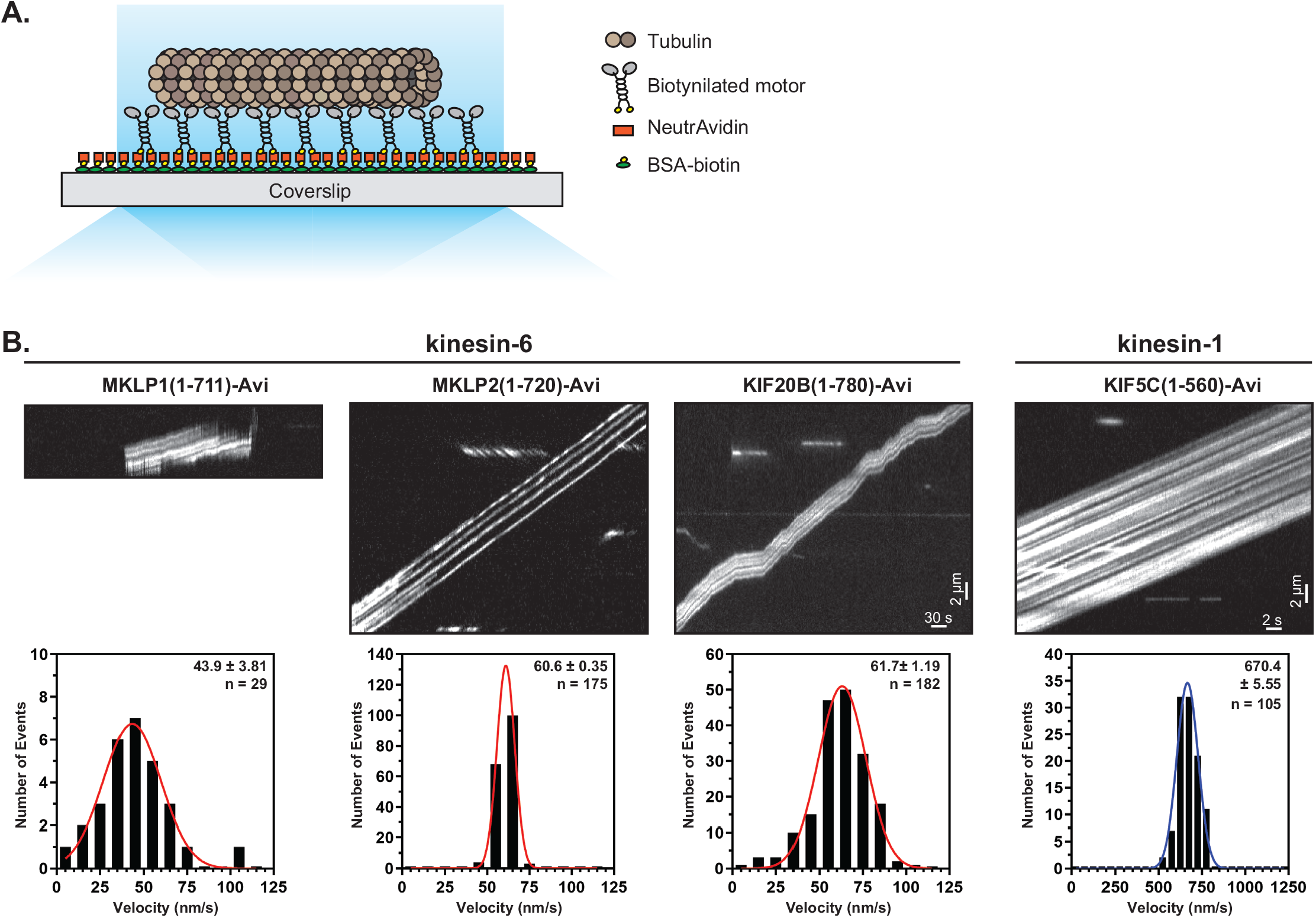
Multimotor properties of kinesin-6 motors in microtubule gliding assays. (A) Schematic of the microtubule gliding assay. Biotinylated kinesin motors were bound to NeutrAvidin-coated coverslips and motor-driven microtubule gliding was imaged using TIRF microscopy. (B) Representative kymographs of the indicated kinesin-6 motors imaged at 1 frame/2 s and the kinesin-1 control imaged at 1 frame/100 ms. Time is on the x-axis and distance is on y-axis. The velocity of microtubule gliding was calculated from kymographs and plotted as a histogram for the population. Insets display mean ± SEM and number of gliding events across four independent experiments for each construct.

### Kinesin-6 proteins can work in teams to drive organelle transport in cells

To test whether kinesin-6 proteins can work in teams to drive the transport of membrane-bound cargoes in cells, we utilized a cargo dispersion assay in which motor proteins are targeted to organelles in a rapamycin-inducible manner and the resulting dispersion of that organelle is utilized as a measurement of the motor’s ability to drive cargo transport (Figure 4A) [78]. For this assay, the kinesin-6 proteins were tagged with mNeonGreen (mNG) and a rapamycin binding domain (FRB). As a control, we characterized the localization of the expressed kinesin-6 proteins and the control kinesin-1 KIF5C(1-560) in COS-7 cells, whose large, flat morphology makes them preferred for imaging changes in organelle localization by fluorescence imaging. For MKLP1, the truncated MKLP1(1-711)-mNG-FRB and full-length MKLP1(1-856)-mNG-FRB proteins expressed at only low levels in the majority of cells and localized to the midbody of one daughter cell after cell division (Figure 3A, red boxes). In cells with higher levels of expression, both MKLP1 versions localized along microtubules throughout the cytosol (Figure 3A). For KIF20B, the truncated KIF20B(1-780)-mNG-FRB and KIF20B(1-982)-mNG-FRB proteins also expressed at only very low levels in the majority of cells and localized to the midbody, however, the KIF20B proteins were found on both sides of the midbody (Figure 3C, red boxes). At higher levels of expression, the KIF20B proteins localized diffusely throughout the cell and not along interphase microtubules (Figure 3C). For MKLP2, the truncated MKLP2(1-720)-mNG-FRB and MKLP(1-770)-mNG-FRB proteins localized diffusely throughout the cell, with faint localization to interphase microtubules in some cells, and did not persist at the midbody after completion of cell division (Figure 3B).

**Figure 3.**
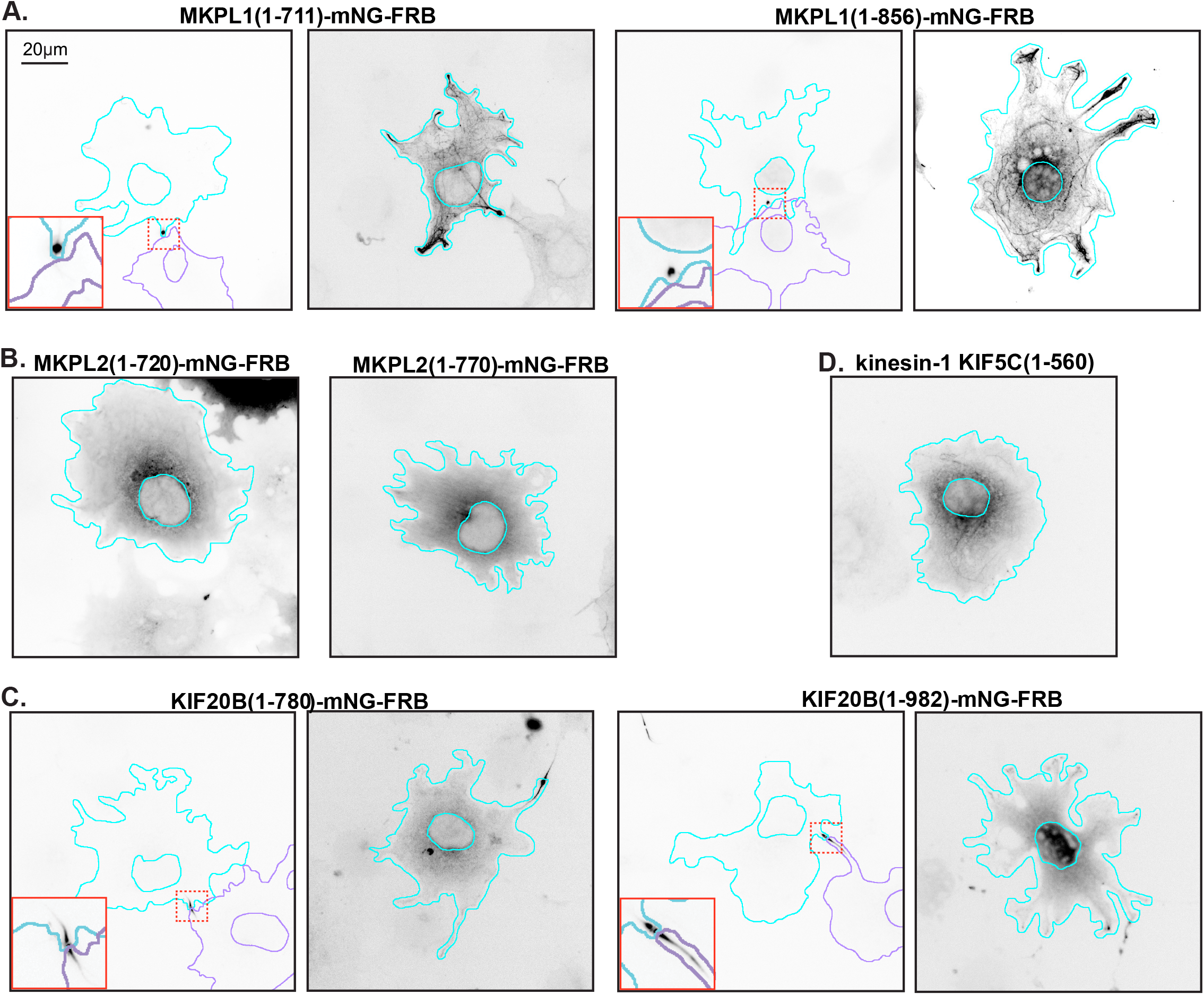
Localization of kinesin-6 motor constructs in interphase cells. Representative images of the localization of (A) truncated MKLP1(1-711) and full-length MKLP1(1-856), (B) truncated MKLP2(1-720) and MKLP2(1-770), (C) truncated KIF20B(1-780) and (1-982), and (D) kinesin-1 KIF5C(1-560) in COS-7 cells. Images are displayed in inverted grayscale. Cyan lines and purple lines indicate the nucleus and periphery of a transfected cell. Red boxes in the lower left corner show a magnified view of the midbody region indicated by the boxes with dotted red lines. Scale bar, 20 μm.

**Figure 4.**
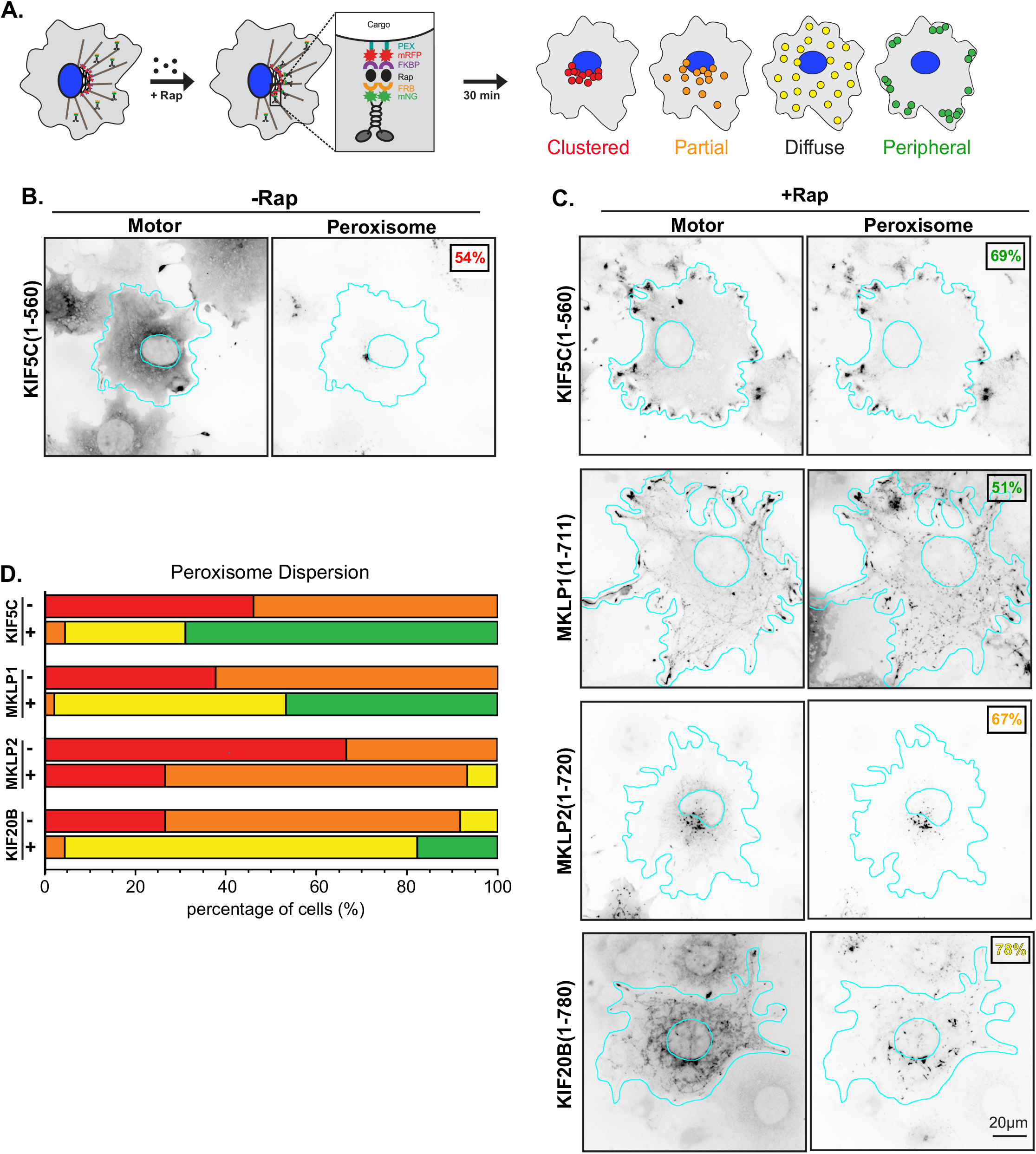
Transport of low-load cargo (peroxisomes) by teams of kinesin-6 motors in cells. (A) Schematic of the inducible cargo dispersion assay. COS-7 cells express a motor tagged with mNG and FRB domain (motor-mNG-FRB) together with a peroxisome-targeting sequence tagged with mRFP and FKBP domain (PEX-mRFP-FKBP). Rapamycin (Rap) causes heterodimerization of the FRB and FKBP domains and recruitment of motors to the cargo membrane. Peroxisome localization in the absence and presence of Rap was qualitatively scored as: red, clustered; orange, partially dispersed; yellow, diffusely dispersed; or green, peripheral. (B,C) Representative images of motor-mNG-FRB and PEX-mRFP-FKBP localization in the (B) absence of Rap or (C) 30 min after addition of Rap. Images are displayed in inverted grayscale. For each condition, the image displays the most frequent phenotype as indicated by the colored number in the upper corner. Cyan lines: nucleus and periphery of each cell. Scale bar, 20 μm. (D) Peroxisome dispersion data for each construct plotted as a stacked bar plot. N>45 cells from three experiments for each construct.

We next tested whether teams of MKLP1, MKLP2, or KIF20B motors could drive transport of membrane-bound cargoes in cells. We first tested whether the kinesin-6 motors could work in teams to drive dispersion of peroxisomes in cells. Peroxisomes localize to the perinuclear region of COS-7 cells, and are relatively immotile under natural conditions [79, 80]. We consider the peroxisome to be a low-load cargo, as it takes about 2-12 pN to move it from its natural location [81, 82]. Motor-mNG-FRB constructs were co-expressed with a peroxisome-targeted FKBP protein tagged with mRFP (PEX-RFP-FKBP). The addition of rapamycin causes dimerization of the FRB and FKBP domains, resulting in targeting of the motor protein to the peroxisome (Figure 4A). Peroxisome localization at 0 and 30 minutes after addition of rapamycin was examined in fixed cells by fluorescence microscopy (Figure 4B,C) and qualitatively scored as clustered (no motor-driven dispersion), partial dispersion, diffuse dispersion, and peripheral (complete peroxisome dispersion to the cell periphery) (Figure 4A). We found that in cells where MKLP1- or KIF20B-mNG-FRB proteins localized to the midbody (Figure 3A,C), the motor could not be recruited to the peroxisome surface upon addition of rapamycin. Thus, these cells were omitted from analysis, and only cells in which the motor was targeted to the peroxisome were scored for dispersion.

MKLP1(1-711)-mNG-FRB was able to drive robust peroxisome dispersion with 97.8% of cells showing peroxisomes with diffuse or peripheral localization (Figure 4C,D), compared to 0% without rapamycin (Figure 4D). This transport ability is comparable to that of the control kinesin-1 KIF5C(1-560) protein (Figure 4C,D). Similarly, KIF20B(1-780)-mNG-FRB was also able to drive peroxisome dispersion as 95.6% of cells displayed diffuse or peripheral peroxisome localization after rapamycin (Figure 4C,D), compared to 8.3% without rapamycin (Figure 4D). However, MKLP2(1-720)-mNG-FRB was largely unable to drive peroxisome dispersion as the majority of cells (66.7%) showed only a partial dispersion and only 6.7% of cells showed diffuse peroxisome localization (Figure 4C,D).

To test the ability of kinesin-6 motors to work in teams and produce force required for cargo transport in cells, we carried out the same assay, but using the Golgi apparatus as the cargo. The Golgi is also localized to the perinuclear region of COS-7 cells and its localization is maintained by a variety of mechanisms including microtubule minus-end directed activity of dynein and is further tethered by myosin motors and linker proteins [83]. We thus consider the Golgi to be a high-load cargo and recent research suggests that it takes ∼200 pN of force to be dispersed from its perinuclear position [84].

Kinesin-6 motors tagged with mNG and FRB were coexpressed with a Golgi-targeted FKBP protein (GMAP210-RFP-FKBP). Golgi localization was examined at 0 and 30 min after the addition of rapamycin and the same categorical analysis was used to quantify motor-driven dispersion of the Golgi as a cargo (Figure 5C). MKLP1(1-711)-mNG-FRB was again able to drive robust transport of Golgi cargo, with 97.8% of cells displaying Golgi localized as either diffuse or peripheral (Figure 5B,C) compared to 0% without rapamycin (Figure 5C). KIF20B(1-780)-mNG-FRB also showed strong dispersion in this assay with 96.9% of cells displaying Golgi localized within the diffuse or peripheral categories (Figure 5B,C), as compared to 0% without rapamycin (Figure 5C). Thus, both MKLP1 and KIF2B show cargo transport properties comparable to that of the control kinesin-1 KIF5C(1-560) protein (Figure 5B,C). Similar to what we observed for the dispersion of low-load peroxisomes, MKLP2(1-720)-mNG-FRB was unable to drive transport of high-load Golgi elements as cells expressing this motor showed either diffuse or peripheral dispersion in only 5% of cells and a partial dispersion in only 37.5% of cells (Figure 5B,C). Taken together, the cargo-dispersion assays suggest that MKLP1 and KIF20B, but not MKLP2, can work in teams to drive transport of membrane-bound cargoes in cells.

**Figure 5.**
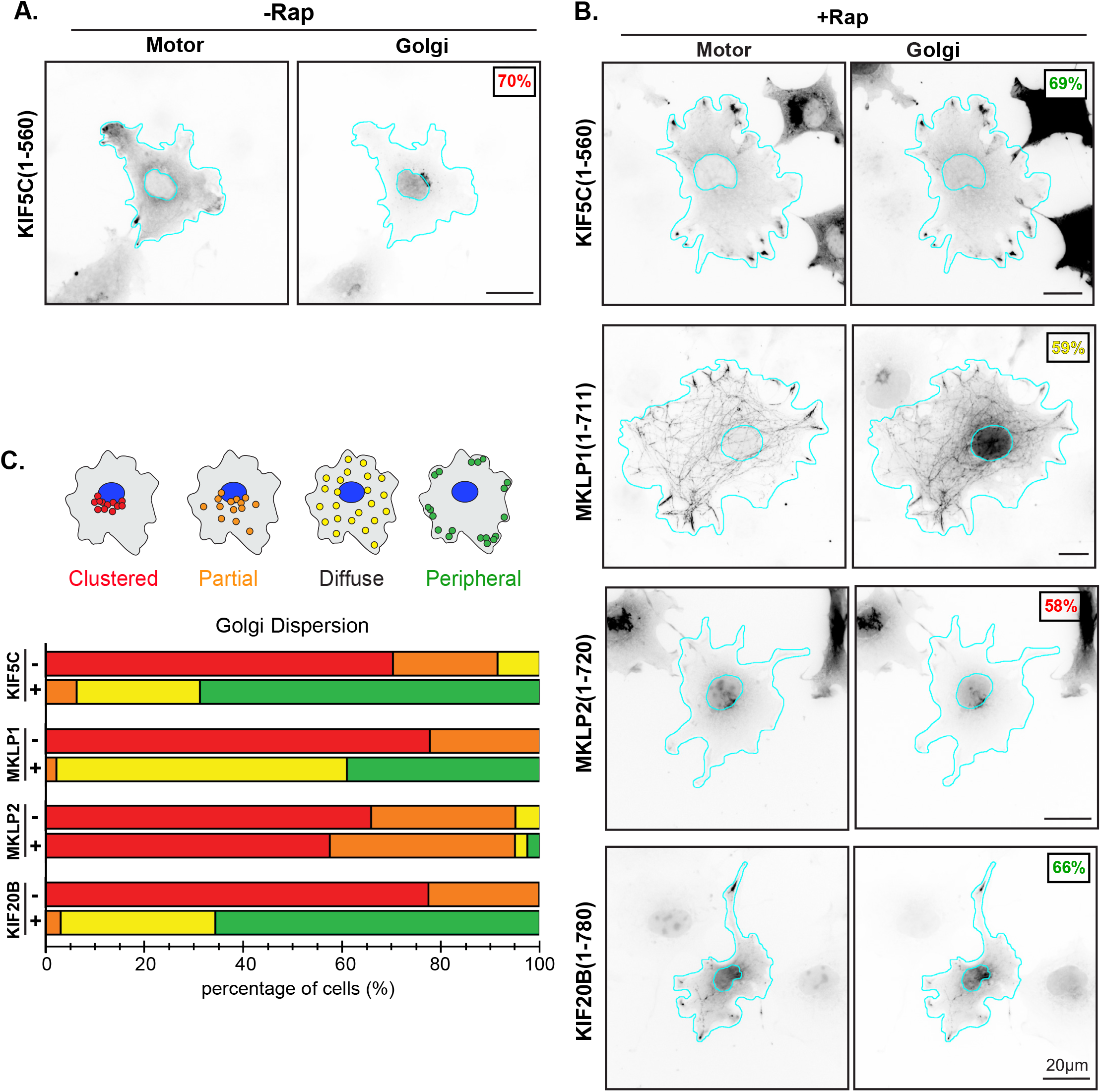
Transport of high-load cargo (Golgi elements) by teams of kinesin-6 motors in cells. (A,B) Representative images of motor-mNG-FRB and Golgi-mRFP-FKBP protein localization in the (A) absence of Rap or (B) 30 min after addition of Rap. Images are displayed in inverted grayscale. For each condition, the image displays the most frequent phenotype as indicated by the colored number in the upper corner. Cyan lines: nucleus and periphery of each cell. Scale bar, 20 μm. (C) Golgi localization in individual cells was scored as clustered (red), partially dispersed (orange), diffusely dispersed (yellow), or peripheral (green). The data for each construct is plotted as a stacked bar plot. N>45 cells from three independent experiments for each construct.

## Discussion

### Kinesin-6 proteins are largely non-processive as single motors

Prior research on the kinesin-6 family of motors has left an incomplete picture of their motility properties. Here, we used single-molecule motility assays as well as team-based cargo transport assays to begin to understand these motor’s underlying motility properties. We find that for mammalian members of the kinesin-6 family, the majority of proteins do not undergo unidirectional processive motility as single molecules but rather diffuse along the surface of the microtubule for several seconds (Figure 1). We also observed, however, that a fraction of motors [most prevalent for MKLP2(1-720) at 23% of events, Figure 1] are able to undergo processive motility at slow speeds. For MKLP1, our results are consistent with previous work showing that a truncated version of the *C*.*elegans* homolog, ZEN-4(1-585), does not display unidirectional processive motility [62]. For MKLP2, the fact that this motor can undergo slow processive motility is consistent with the work of Andriaans et al with full-length MKLP2-EGFP, although these authors did not report whether any of the molecules undergo diffusive rather than processive motion [29].

It was previously suggested that the presence of an extended region (∼60 aa) that separates the kinesin-6 neck linker and neck coil regions (Figure S1A) would prevent coordination of the two motor domains and processive motility [12, 18, 27, 53, 57]. Indeed, kinesin proteins with extended neck linker domains have been found to interact statically with microtubules [85-88]. However, recent work showed that the plant Phragmoplast-associated kinesin-related protein 2 (PAKRP2) exhibits processive unidirectional motility on microtubules as individual homodimers [89]. These results indicate that the relationship between neck linker length and kinesin stepping is not as straight-forward as previously thought. Whether the non-processive nature of the kinesin-6 motors is additionally impacted by the N-terminal extension and/or the insertion in loop-6 remains to be determined, however, structural work indicates that the loop-6 insertion does not impact the microtubule-binding or catalytic activity of *Ce*ZEN-4 [52, 54].

It is presently unclear why only a subset of kinesin-6 molecules are capable of directional motility (Figure 1 and [29]) but we note that similar results have been observed in cells where MKLP2-CPC cargo complexes showed largely diffusive motion on the central spindle with only a fraction of complexes undergoing processive motility [27, 29]. One possibility is that processive motility results from the clustering of dimeric non-processive motors. For example, *Ce*ZEN-4 can naturally oligomerize and form higher-order clusters that undergo slow processive motility as single particles [62]. We were unable to observe oligomerization of our MKLP1(1-711) or MKLP1(1-856) constructs perhaps because the clustering element of *Ce*ZEN-4 (aa 585-601, [62]) is not conserved in *Hs*MKLP1. Clustering of MKLP1 in our system may also be hindered by the low expression and thus protein concentration of MKLP1 in mammalian cells and/or the requirement for other cellular factors, for example MgcRacGAP, Aurora B, and 14-3-3 proteins [18, 32, 57, 63], for clustering. An alternative possibility is that kinesin-6 motors can bind to the microtubule lattice but these interactions can only infrequently be converted into productive interactions that lead to motility.

### Kinesin-6 motors can work in teams for force generation and cargo transport

An inability of single kinesin proteins to undergo unidirectional processive motility does not preclude their ability to work in teams to drive transport [58-61]. Indeed, we find that that all of the mammalian kinesin-6 motors can work in teams to drive microtubule gliding at slow speeds, with mean velocities of 43.9 nm/s for MKLP1, 60.6 nm/s for MKLP2, and 61.7 nm/s for KIF20B (Figure 2). These speeds are consistent with previous work using gliding assay to investigate the motility of MKLP1 and its homologs *Ce*ZEN-4 and *Dm*Pavarotti as well as KIF20B [14, 18, 49, 57, 62-64]

Microtubule gliding assays utilize kinesin motors stably attached to a surface and thus cannot predict how motors will cooperate when attached to and diffusing within a lipid bilayer. Indeed, we find that kinesin-6 motors differ in their ability to drive transport of membrane-bound cargoes in cells. MKLP1 and KIF20B can work effectively in teams to drive the dispersion of both low-load and high-load membrane-bound organelles in cells, suggesting that these motors may be capable of high force output. In contrast, MKLP2 was only able to drive the dispersion of low-load peroxisomes in the same assay (Figures 4,5), suggesting that the force output of this motor may be impaired relative to kinesin-1.

Force generation by kinesin motors requires neck linker docking which occurs in two sequential steps: zippering of the neck linker with the coverstrand to form the cover-neck bundle (CNB) followed by latching of the neck linker to the surface of the motor domain via a conserved asparagine residue (N-latch) ([55, 56] Figure S1D). Of the kinesin-6 motors, KIF20B is the only member that contains the N-latch residue (Figure S1D) and is thus predicted to be capable of high force production. In support of this prediction, we find that KIF20B can robustly drive transport of both low-load peroxisomes and high-load Golgi in cells (Figures 4,5) and previous work implicated KIF20B in transport of Shootin to the end of the axon [90].

MKLP1 and MKLP2 lack the N-latch residue [D in MKLP1 (HsKIF23) and Q in MKLP2 (HsKIF20A), Figure S1D], suggesting that their force output may be lower than that of kinesin-1. However, our results show that MKLP1 is fairly efficient at both low-load and high-load transport in cells (Figures 4 & 5). Thus, the presence of the N-latch residue does not correlate with the ability to generate a high force output and thus the force-generation mechanism of MKLP1 is unclear.

In contrast, our results with MKLP2 are consistent with the hypothesis that this motor lacks key residues for kinesin force generation as MKLP2 is unable to efficiently drive the transport of either low-load or high-load cargoes to the cell periphery (Figures 4 & 5). Interestingly, in addition to lacking the N-latch residue, the neck linker of MKLP2 contains a number of glycine and proline substitutions (Figure S1D) that have a very low propensity to adopt a β-sheet formation [91, 92]. These substitutions could result in impaired neck linker docking and indeed, Atherton et al. were unable to resolve a docked NL conformation for MKLP2 even in the ATP- and microtubule-bound state [53]. Previous work suggested that MKLP2 could function as a transporter to carry CPC substrates towards the cell equator as MKLP2 rigor mutants concentrate at the spindle poles [29, 93]. Our finding that MKLP2 is only capable of low-load transport implies that transport of CPC complexes does not require a high force output by the transporting motor.

### Localization in interphase cells

Recent work suggests that kinesin-6 motors have functional roles in interphase of cycling cells as well as in post-mitotic cells. We find that in the majority of interphase cultured cells, truncated (1-711) and full-length (1-856) constructs of MKLP1 accumulate on one side of the midbody. The asymmetric distribution of MKLP1 is consistent with previous work showing that MKLP1 is a marker of the midbody ring remnant that is asymmetrically inherited by one daughter cell after completion of abscission [12, 94-96] and can be detected extracellularly secreted midbody remnants [97]. In cells with higher expression of MKLP1(1-711) and MKLP1(1-856), the proteins localized along the length of cytosolic microtubules, consistent with previous work analyzing the localization of MKLP1 tail-deletion mutants in interphase HEK293 cells [70]. In contrast to the asymmetric localization of MKLP1, the expressed KIF20B proteins associate with both sides of the midbody on the microtubule ‘arms’. That KIF20B persists at this location into interphase (Figure 3) is consistent with previous work [51, 90] whereas MKLP2 is lost after furrow ingression is complete [98].

## Material and Methods

### Plasmids

A truncated, constitutively active kinesin-1 [rat KIF5C(1-560)] was used as a control in all experiments [99]. Plasmids containing cDNAs encoding the human kinesin-6 family members MKLP1 isoform 2 (*Hs*KIF23, Uniprot Q02241-2), MKLP2 isoform 1 (*Hs*KIF20A, Uniprot O95235, gift of Ryoma Ohi [27]), and *Hs*KIF20B Isoform 3 (Uniprot Q96Q89-3, gift of Orly Reiner [90]). The truncated versions MKLP1(1-711), MKLP2(1-720), and KIF20B(1-780) were generated by a combination of PCR, Gibson cloning, and gene synthesis. All plasmids were verified by DNA sequencing. MKLP1(1-711) lacks the insert present in KIF23 isoform 1 (also known as CHO1, Uniprot Q02241-1 [16]) and thus likely reflects the core motor properties of both CHO1 and MKLP1 isoforms. KIF20B contained the protein sequence conflict E713K and natural variations N716I and H749L. Motors were tagged with three tandem monomeric Citrine fluorescent proteins (3xmCit) for single-molecule imaging assays [77] or monomeric NeonGreen (mNG)-FRB for inducible cargo-dispersion assays in cells. The peroxisome-targeting PEX3-mRFP-FKBP construct was a gift from Casper Hoogenraad [78]. The Golgi-targeting GMAP210p-mRFP-FKBP construct is described in [100]. Constructs coding for FRB (DmrA) and FKBP (DmrC) sequences were obtained from ARIAD Pharmaceuticals and are now available from Takara Bio Inc. Plasmids encoding monomeric NeonGreen were obtained from Allele Biotechnology.

### Cell culture, transfection, and lysate preparation

COS-7 (African green monkey kidney fibroblasts, American Type Culture Collection, RRID:CVCL_0224) were grown at 37°C with 5% (vol/vol) CO_2_ in Dulbecco’s Modified Eagle Medium (Gibco) supplemented with 10% (vol/vol) Fetal Clone III (HyClone) and 2 mM GlutaMAX (L-alanyl-L-glutamine dipeptide in 0.85% NaCl, Gibco). Cells are checked annually for mycoplasma contamination and were authenticated through mass spectrometry (the protein sequences exactly match those in the African green monkey genome). 24 hr after seeding, the cells were transfected with plasmids using TransIT-LT1 transfection reagent (Mirus), and Opti-MEM Reduced Serum Medium (Gibco). Cells were trypsinized and harvested 24 hr after transfection by low-speed centrifugation at 3000 x *g* at 4°C for 3 min. The pellet was resuspended in cold 1X PBS, centrifuged at 3000 x *g* at 4°C for 3 min, and the pellet was resuspended in 50 μL of cold lysis buffer [25 mM HEPES/KOH, 115 mM potassium acetate, 5 mM sodium acetate, 5 mM MgCl_2_, 0.5 mM EGTA, and 1% (vol/vol) Triton X-100, pH 7.4] with 1 mM ATP, 1 mM phenylmethylsulfonyl fluoride, and 1% (vol/vol) protease inhibitor cocktail (P8340, Sigma-Aldrich). Lysates were clarified by centrifugation at 20,000 x *g* at 4°C for 10 min and lysates were snap frozen in liquid nitrogen and stored at −80°C.

### Single-Molecule Motility Assays

Microtubules were polymerized (purified porcine tubulin unlabeled and HiLyte-647-labeled, Cytoskeleton Inc) in BRB80 buffer [80 mM Pipes/KOH pH 6.8, 1 mM MgCl_2_, 1 mM EGTA] supplemented with GTP and MgCl_2_ and incubated for 60 min at 37°C. 20 μM taxol in prewarmed BRB80 was added and incubated for 60 min to stabilize microtubules. Microtubules were stored in the dark at room temperature for up to 2 weeks. Flow cells were prepared by attaching a #1.5 18 mm^2^ coverslip (Thermo Fisher Scientific) to a glass slide (Thermo Fisher Scientific) using double-sided tape. Microtubules were diluted in fresh BRB80 buffer supplemented with 20 μM taxol, infused into flow cells, and incubated for five minutes to allow for nonspecific absorption to the glass. Flow cells were then incubated with blocking buffer [5 mg/mL casein in P12 buffer supplemented with 5 μM taxol] for five minutes. Flow cells were then infused with motility mixture [0.5–5.0 μL of COS-7 cell lysate, 25 μL BRB80 buffer, 1 μL 100 mM ATP, 0.5 μL 100 mM MgCl_2,_ 0.5 μL 100 mM DTT, 0.5 μL 20 mg/mL glucose oxidase, 0.5 μL 8 mg/mL catalase, and 0.5 μL 1 M glucose], sealed with molten paraffin wax, and imaged on an inverted Nikon Ti-E/B total internal reflection fluorescence (TIRF) microscope with a perfect focus system, a 100 × 1.49 NA oil immersion TIRF objective, three 20 mW diode lasers (488 nm, 561 nm, and 640 nm) and EMCCD camera (iXon^+^ DU879; Andor). Image acquisition was controlled using Nikon Elements software at 1 frame/50 ms or 1 frame/2 s and all assays were performed at room temperature. Motility data were analyzed by first generating maximum intensity projections to identify microtubule tracks (width = 3 pixels) and then generating kymographs in Fiji/ImageJ (National Institutes of Health). Events that ended as a result of a motor reaching the end of a microtubule were included; therefore, the reported dwell times may be an underestimation. For each motor construct, GraphPad Prism was used to bin the dwell times and generate a histogram by plotting the number of motility events for each bin.

### Protein Purification

MKLP1(1-711)-3xFLAG-Avi was cloned by stitching four oligonucleotide primer sequences together into a digested MKLP1(1-711)-Avitag plasmid using a HiFi DNA assembly reaction kit (M5520A, New England Biolabs). Ten 10cm plates were seeded with COS-7 cells and each plate was co-transfected 24 hours later with 4.08 μg MKLP1(1-711)-3xFLAG-Avi and 4.08 μg HA-BirA plasmids using TransIT-LT1 transfection reagent (Mirus), and Opti-MEM Reduced Serum Medium (Gibco). Cells were trypsinized and harvested 24 hr after transfection by low-speed centrifugation at 3000 x *g* at 4**°**C for 3 min. The pellet was resuspended in cold 1X PBS, centrifuged at 3000 x *g* at 4**°**C for 5 min, and the pellet was resuspended in 1000 μL of cold lysis buffer with 1 mM ATP, 1 mM phenylmethylsulfonyl fluoride, 1 mM DTT and 1% (vol/vol) protease inhibitor cocktail (P8340, Sigma-Aldrich). Cells were centrifuged at 20,000 x *g* at 4**°**C for 10 min and the pellet was discarded. 50 μL anti-FLAG M2 beads (A2220, Sigma-Aldrich) were mixed into the supernatant for 1.5 hours then washed twice with 2xFLAG wash buffer [300 mM KCl, 40 mM Imidazole/HCl, 10 mM MgCl_2_, 2 mM EDTA, and 2mM EGTA] supplemented with 1 mM phenylmethylsulfonyl fluoride, 1 mM DTT, and 1% (vol/vol) protease inhibitor cocktail (P8340, Sigma-Aldrich). 3 mM ATP was added for the first wash only. Protein was eluted from the beads with 100 μL elution buffer [1% (vol/vol) BRB80, 1% (vol/vol) protease inhibitor cocktail (P8340, Sigma-Aldrich), 1 mM phenylmethylsulfonyl fluoride, 0.5 mM DTT, 0.1 mM ATP, and 0.5 mg/ml FLAG peptide (F4799, Sigma-Aldrich)]. The beads were pelleted by centrifugation at 20,000 x *g* at 4**°**C for 10 min and aliquots were snap frozen in liquid nitrogen and stored at −80**°**C.

### Microtubule Gliding Assay

A flow cell was prepared, and microtubules were assembled as described in the single molecule section. Biotinylated motors were generated by coexpression of motors tagged with the 15-aa Avi tag and the bacterial biotin ligase BirA fused with HA tag (HA-BirA) in COS-7 cells. Biotinylated motors were attached to the coverslip surface by sequential incubation of flow cells with (A) 1 mg/ml BSA-biotin, (B) blocking buffer [0.5 mg/ml casein and 10 µM taxol in BRB80], (C) 0.5 mg/ml NeutrAvidin, (D) blocking buffer, and (E) cell lysates with 2 mM ATP, 10 mg/ml casein in BRB80, 10 μM taxol in BRB80, and blocking buffer. Taxol-stabilized HiLyte 647–labeled microtubules in motility mixture [2 mM ATP, 10 µM taxol, 2 mM MgCl_2_, and oxygen scavenging in BRB80] were then added, and the flow cells were sealed with molten paraffin wax and imaged by TIRF microscopy. For KIF5C, images were acquired continuously at 50 ms per frame for 30 s. For MKLP1, MKLP2, and KIF20B, images were acquired at one frame every 2 s for 10 min. Maximum-intensity projections were generated, and the kymographs were produced by drawing along these tracks (width = 3 pixels) using ImageJ. Stalled events were ignored. Velocity was defined as the distance on the y axis of the kymograph divided by the time on the x axis of the kymograph.

### Inducible Cargo Dispersion Assays

Plasmids for expression of kinesin-1 or any of the kinesin-6 family motors tagged with monomeric NeonGreen and an FRB domain were cotransfected into COS-7 cells with a plasmid for expression of PEX3-mRFP-FKBP or GMAP210p-mRFP-2xFKBP at a ratio of 5:1 with TransIT-LT1 transfection reagent (Mirus). Eighteen hours after transfection, rapamycin (Calbiochem, Sigma) was added to final concentration of 44 nM to promote FRB and FKBP heterodimerization and recruitment of motor the peroxisome or Golgi surface. Cells were fixed with 3.7% formaldehyde (Thermo Fisher Scientific) in 1X PBS, quenched in 50 mM ammonium chloride in PBS for 5 min, permeabilized for 5 min in 0.2% Triton-X 100 in PBS for 5 min and blocked in 0.2% fish skin gelatin in PBS for 5 min. Primary and secondary antibodies were added to blocking buffer and incubated for 1 hr at room temperature. Primary antibodies: polyclonal antibody against cis-Golgi marker giantin (1:1200 PRB-114C, Covance). Secondary antibodies: goat anti-rabbit Alexa680-labeled secondary antibody (1:500, Jackson ImmunoResearch). Cell nuclei were stained with 10.9 μM 4′,6-diamidino-2-phenylindole (DAPI, 1:1000 Sigma). Coverslips were mounted in ProlongGold (Invitrogen) and imaged using an inverted epifluorescence microscope (Nikon TE2000E) with a 40 × 0.75 NA objective and a CoolSnapHQ camera (Photometrics). Only cells expressing low levels of motor-mNG-FRB were imaged and included in quantification, as high expression of KIF5C disrupted the microtubule network. Cargo localization before and after motor recruitment was qualitatively assessed. The phenotype of cargo dispersion was scored as clustered, partial, diffuse, or peripheral dispersion based on the signal localization in the PEX3 (peroxisome) or GMAP210p (Golgi) signal.

## Acknowledgements

We are grateful to Kristin Schimert, Yang Yue, Lynne Blasius and other members of the Verhey laboratory for advice, support, discussions, and reagents. This work was supported by grants from the National Institutes of Health to KJV (R01GM070862, R35GM131744). BGB was supported by the Cellular and Molecular Biology Training Grant T32-GM007315 from the National Institutes of Health, a Graduate Research Fellowship (DGE 1256260) from the National Science Foundation, an EDGE Fellowship from the Endowment of Basic Sciences at the University of Michigan Medical School, and a Rackham Predoctoral Fellowship from the Horace H. Rackham School of Graduate Studies at the University of Michigan. AP was supported by the Undergraduate Research Opportunities Program (UROP) at the University of Michigan and the Undergraduate Honors Program in Molecular, Cellular, and Developmental Biology.

## Author Contributions

Conceptualization: all authors

Investigation and Analysis: A.P.

Writing – Original Draft: A.P.

Writing – Review & Editing: all authors

Funding Acquisition and Resources: K.J.V

Project Administration: B.G.B. and K.J.V.

---

**Figure S1.**
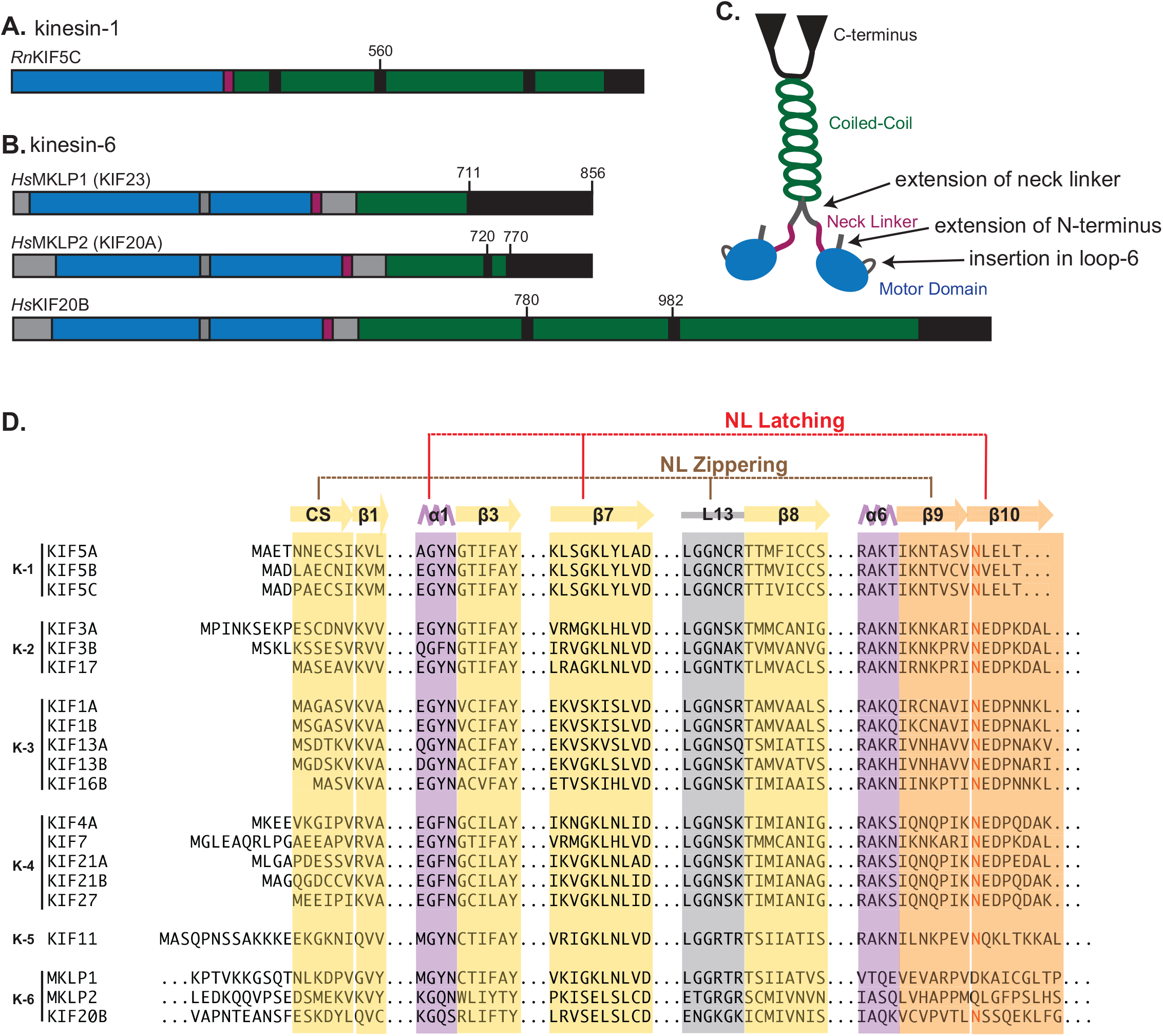
Domain organization and unique sequence features of kinesin-6 motors. (A,B) Domain organization of (A) kinesin-1 and (B) kinesin-6 motor proteins. Blue, motor domain; green, coiled-coil; purple, neck linker; gray, insertions in kinesin-6 relative to kinesin-1. Numbers and black lines indicate the positions of protein truncations. (C) Schematic. Kinesin-6 motor proteins are defined by three sequence features: an extension of the N-terminus before the core kinesin motor domain, an insertion in loop-6 on the surface of the core kinesin motor domain, and an extended sequence between the neck linker and the neck coil. (C) Alignment of sequences critical for neck linker docking in kinesin motor proteins. Neck linker docking occurs in two sequential steps. First, NL zippering which entails zippering of β9 of the neck linker with the coverstrand (CS) to form the cover-neck bundle (CNB). Second, NL latching which entails interactions of β10 of the neck linker with β7 and α1 regions and latching via a conserved asparagine residue (N-latch, red text).

**Figure S2.**
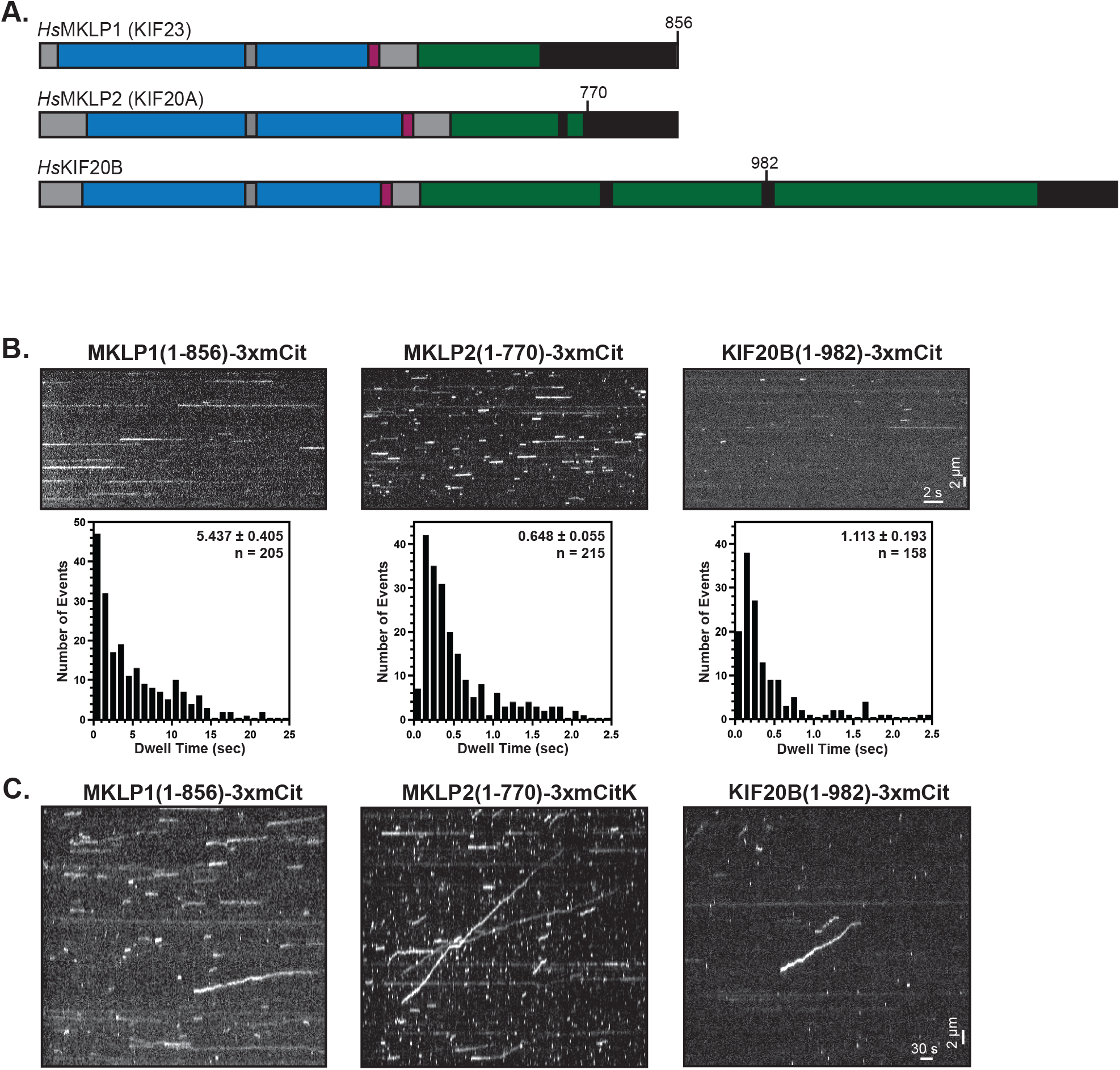
Single-molecule motility properties of additional kinesin-6 constructs. (A) Domain organization and schematic of kinesin-6 motor proteins. Blue, motor domain; green, coiled-coil; purple, neck linker. Numbers and black lines indicate the positions of protein truncations. (B) Motility properties of the indicated kinesin-6 proteins and the kinesin-1 control using TIRF microscopy and a fast acquisition rate (1 frame/50 ms). Representative kymographs are shown with time on the x-axis (scale bar, 2s) and distance on y-axis (scale bar, 2 μm). Each motor’s dwell time on the microtubule was calculated from kymographs and plotted as a histogram for the population. Insets display the mean ± SEM and number of events across three independent experiments for each construct. (C) Motility properties of indicated kinesin-6 proteins or the kinesin-1 control using TIRF microscopy and a slow acquisition rate (1 frame/2s). Representative kymographs are shown with time on the x-axis (scale bar, 30s) and distance on the y-axis (scale bar, 2 μm).

**Figure S3.**
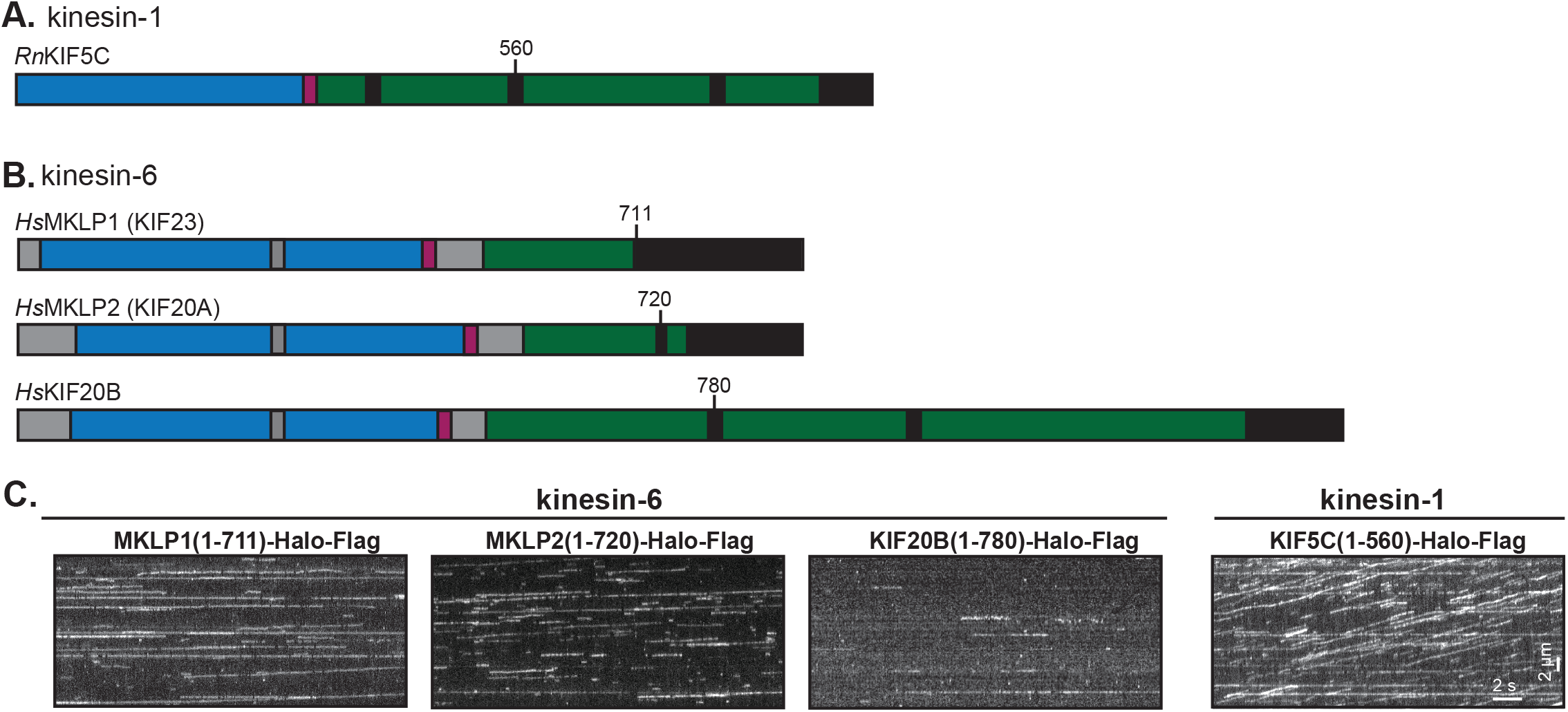
Single-molecule properties of kinesin-6 truncations tagged with Halo tag. (A) Domain organization and schematic of kinesin-6 and kinesin-1 truncations. Blue, motor domain; green, coiled-coil; purple, neck linker. Numbers and black lines indicate the positions of protein truncations. (B) Motility properties of the indicated kinesin-6 proteins and the kinesin-1 control using TIRF microscopy and a fast acquisition rate (1 frame/50 ms). Representative kymographs are shown with time on the x-axis (scale bar, 2s) and distance on y-axis (scale bar, 2 μm).

